# ERROR MODELLED GENE EXPRESSION ANALYSIS (EMOGEA) PROVIDES A SUPERIOR OVERVIEW OF TIME COURSE RNA-SEQ MEASUREMENTS AND LOW COUNT GENE EXPRESSION

**DOI:** 10.1101/2022.02.18.481000

**Authors:** Tobias K. Karakach, Federico Taverna, Jasmine Barra

**Affiliations:** Laboratory of Integrative Multi-Omics Research, Department of Pharmacology, Dalhousie University, Halifax, NS, B3H 4R2, CANADA; Children’s Hospital Research Institute of Manitoba, Winnipeg, MB R3E 3P4, CANADA; Department of Pathology, Dalhousie University, Halifax, Nova Scotia, NS, B3H 4R2, CANADA

## Abstract

Serial RNA-seq studies of bulk samples are widespread and provide an opportunity for improved understanding of gene regulation during *e*.*g*., development or response to an incremental dose of a pharmacotherapeutic. In addition, the widely popular single cell RNA-seq (scRNA-seq) data implicitly exhibit serial characteristics because measured gene expression values recapitulate cellular transitions. Unfortunately serial RNA-seq data continue to be analyzed by methods that ignore this ordinal structure and yield results that are difficult to interpret. Here, we present Error Modelled Gene Expression Analysis (EMOGEA), a principled framework for analyzing RNA-seq data that incorporates measurement uncertainty in the analysis, while introducing a special formulation for modelling data that are acquired as a function of time or other continuous variable. By incorporating uncertainties in the analysis, EMOGEA is specifically suited for RNA-seq studies in which low-count transcripts with small fold-changes lead to significant biological effects. Such transcripts include signaling mRNAs and non-coding RNAs (ncRNA) that are known to exhibit low levels of expression. Through this approach, missing values are handled by associating with them disproportionately large uncertainties which makes it particularly useful for single cell RNA-seq data. We demonstrate the utility of this framework by extracting a cascade of gene expression waves from a well-designed RNA-seq study of zebrafish embryogenesis and, a scRNA-seq study of mouse pre-implantation and provide unique biological insights into the regulation of genes in each wave. For non-ordinal measurements, we show that EMOGEA has a much higher rate of true positive calls and a vanishingly small rate for false negative discoveries compared to common approaches. Finally, we provide an R package (https://github.com/itikadi/EMOGEA) that is self-contained and easy to use.

**Graphical Abstract:**
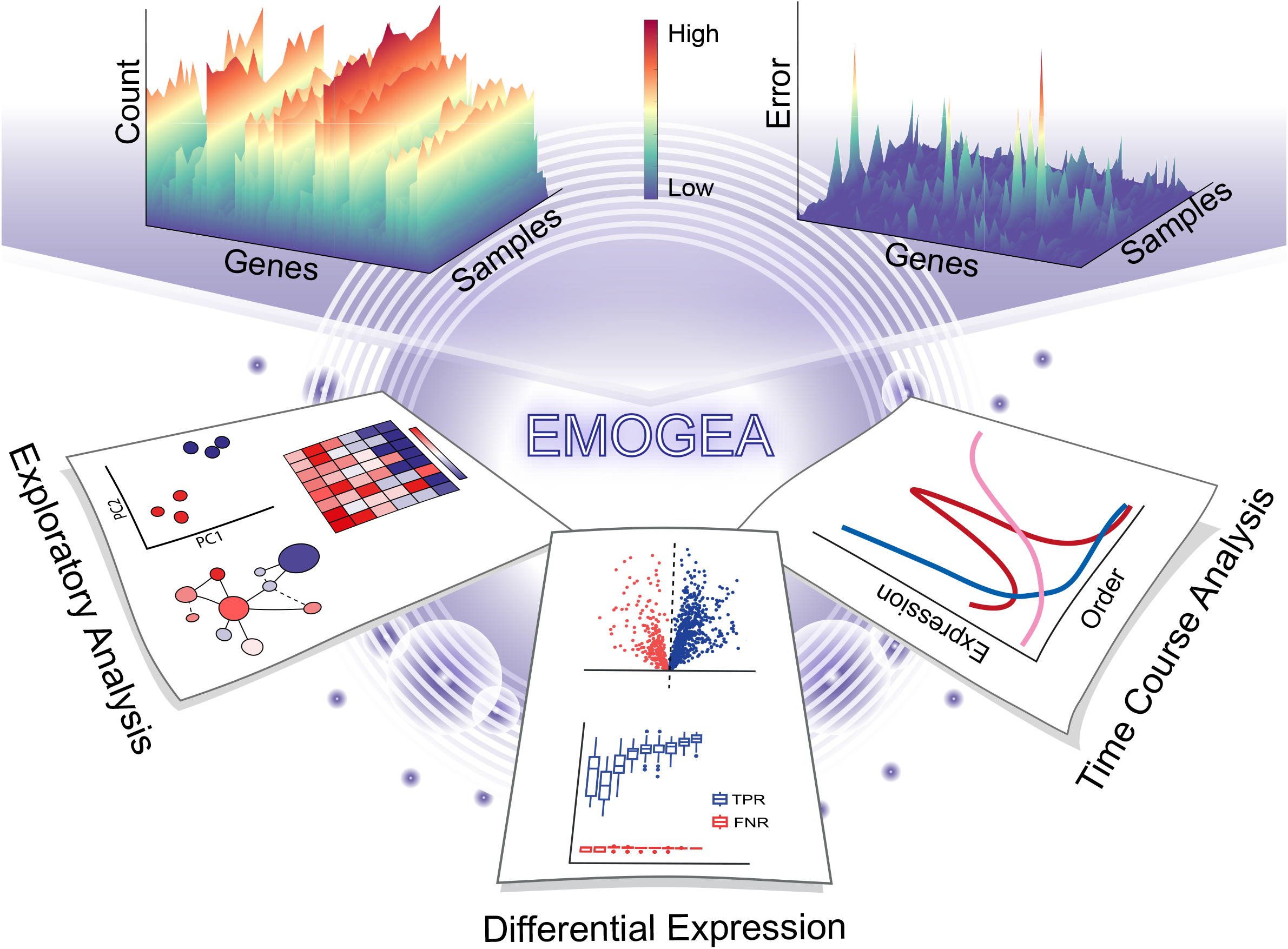
Graphical representation of EMOGEA indicating the incorporation of measurement errors in modeling RNA-seq data to generate superior results in exploratory analysis, differential gene expression analyses and, scRNA-seq and Time Course analyses.

## MAIN TEXT

Next generation sequencing (NGS) has supplanted microarray hybridizations^1^ as a method for the measuring gene expression and identifying differential gene expression (DGE). This is because of both its increased throughput and the larger dynamic range of signals that can be measured. Advances in microfluidics technologies have further enabled NGS to be extended to measuring levels of DNA or RNA in single cells rather than bulk sample homogenates, thereby significantly increasing throughput and resolution^2^. At the inception of microarray technologies, a lot of effort was expended in characterizing the distributional properties of the measurements^3–6^ along with developments of statistical methods to determine DGE^7–10^. When RNA sequencing (RNA-seq) emerged, the same statistical methods were used to analyze the data following transformations that allowed their distributional characteristics to approximately resemble those of their microarray counterparts. Subsequently, other methods were specifically developed to analyze RNA-seq data treating their distributions as Negative Binomial (NB) or over-dispersed Poisson^11,12^ rather than transforming them to mimic microarrays. Despite this better understanding of the distributional characteristics of RNA-seq data, a number of limiting factors persists. First, there is a lack of methods that can incorporate measurement uncertainty in determining DGE^11,13–15^ so as to decrease the rate at which false discoveries are made. Second, time ordered data continue to be analyzed using the classical pairwise comparison approaches that ignore the autocorrelations between successive samples or using splines and other smoothing-based approaches that have limited biological interpretation.

Currently one of the most prominent approaches for handling measurement errors in the analysis of RNA-seq data is to fit a general trend to the NB dispersions^11,12,16^, and use that fit as a surrogate for gene-specific variation. Another alternative is to model gene-wise dispersions through empirical Bayes procedures^16,17^ that borrow information from a set of self-consistent gene expression values under the null hypothesis of no overall differential expression. These approaches do not, unfortunately, address the real question of true measurement uncertainty, relying instead on the intra-sample data spread to infer the over-dispersion parameter.

The impetus for accurately modelling measurement errors in RNA-seq is to produce a reliable list of genes that are differentially expressed and minimize under- or over-estimation. A number of publications^13,18–21^ compare the performance of methods for RNA-seq data analysis using the false discovery rate (FDR) as a standard marker of quality, conditioned on a traditional cut-off of two-fold differential expression. Schurch *et al*.,^21^ have performed one such comprehensive comparison adjusting this two-fold differential gene expression (DGE) cut-off in the context of replication levels. In their findings, it is clear that technical variability plays an important role in FDR control and show that model performance increases with the number of replicates and that the smaller the DGE cut-off the more replicates are needed to control the FDR. In reality there is a finite number of replicates experimentalists can acquire due either to a limit in resources, sample availability or both.

We address these two critical shortcomings in RNA-seq (measurement errors and the analysis of serial data) via the EMOGEA framework that fundamentally shifts the way these data are analyzed by combining techniques for: (a) incorporating measurement error information into the pre-processing steps with, (b) a special approach for modeling ordinal data.

## RESULTS

We assume that measurement errors in RNA-seq arise from well-defined sources that can be quantified through replication. We consider a cumulative response error where the total variance has additive components that derive from these known sources^3,13,22,23^ such that: *σ*^2^_*total*_ = *σ*^2^_*biol*_ + *σ*^2^_*tech*_ + *σ*^2^_*ε*_. In this formulation *σ*^2^_*total*_ represents the total variance, *σ*^2^_*biol*_ the biological variance, *σ*^2^_*tech*_ the experimental (technical) variability while *σ*^2^_*ε*_ is a random component that is assumed to be independent, identical and normally distributed (*iid* normal). *σ*^2^_*tech*_ represents a superset of factors that include batch differences, sample extraction protocols, library preparation, sequencing and computational errors, while *σ*^2^_*biol*_ is the true biological question for which the experiment is designed. We could reduce this formulation into *σ*^2^_*total*_ = *σ*^2^_*biol*_ + *σ*^2^_*samp*_ where the last term includes the smaller random error component that is usually modelled by methods estimating over-dispersion. In the EMOGEA framework it would be clearly absurd to assume that the statistical characteristics *σ*^2^_*samp*_ are *iid* normal given known correlations in gene expression. We demonstrate the utilization of this approach with an application to data acquired under two distinct design strategies.

First, for pairwise comparative design strategies, we begin by determining the magnitude of *σ*^2^_*samp*_. We then incorporate it into the pre-processing step prior to DGEA using limma’s empirical Bayes moderated t-statistic or, exploratory data analyses using principal component analysis (PCA) or multi-dimensional scaling (MDS) after ensuring that the data exhibit log-normal distributional characteristics.

Second, for time-ordered (ordinal) and single cell (sc)RNA-seq experiments, we employ a conceptually simple strategy that uses the principles of bilinear decomposition with non-negativity constraints on the solution, while implicitly incorporating measurement uncertainties in the model development. This approach is referred to as multivariate curve resolution (MCR) and is implemented via weighted alternating least squares. Biologically, the rationale for a bilinear model in ordinal experiments is straightforward and proceeds as follows. Assuming time as the ordinal variable, and given a measurement of gene expression from samples taken successively during this time, we can arrange the data in a matrix form, **X**, such that each row represents a gene while each column an expression value for that gene over time. Using a simple multiple linear regression, we can represent the measured data as:

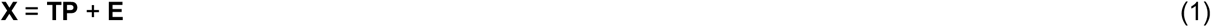

where **E**, is the matrix of random errors while **T** and **P** are smaller matrices that characterize the biological information in **X**. EMOGEA introduces the approach for determining **T** and **P** from RNA-seq data with the constraint that the solution must be non-negative. This constraint is informed by the knowledge that whereas the expression level of a gene can be zero, it can never be negative. A plot of the columns of the matrix **P**, against time indicates how the expression of a class of genes cascades over the course of the experiment while the matrix **T** indicates which genes exhibit the characteristics represented by **P**. A question that arises is how to determine the size of **T** or **P** (corresponding to the number of independent components). There are several established methods for making this determination including examination of standardized residual root mean squares, lack-of-fit measures, Akaike Information Criterion (AIC) among others. Whereas these approaches provide computationally validated solutions, in practice this cut-off for many biological data not clear-cut. In EMOGEA we allow for the flexibility to determine the size of these smaller matrices sequentially by specifying a set number of profiles and examining their graphical characteristics in relation to what is expected biologically.

In practice the only measurements available is the so called “count” matrix **X**. To determine the values of either **T** or **P**, we make an initial guess of either matrix and estimate the other from least squares and impose the non-negativity constraint. We continue this procedure with the second matrix, applying the same constrains and alternate the least squares estimation until a convergence criterion is met. We provide more algorithmic details in the Methods section including the procedure for measurement error incorporation that allows specifying disproportionately large uncertainties for missing values, in effect down-weighting their significance.

We apply EMOGEA to three data types which we will categorize on the basis of the most common design strategies for transcriptomics analyses: Time Course (TC), scRNA-seq and, case-control (CC). CC studies are the most common strategies for transcriptomics studies and involve, in the simplest case, comparing two conditions to determine differential gene expression. In these types of studies, there are two common approaches for visualizing the effect of a treatment: exploratory analysis via methods such as principal component analysis (PCA), multidimensional scaling (MDS) and others or; via analysis of differential expression to identify genes that are expressed at statistically significant levels between the conditions. In contrast, although TC experiments are more informative of the ultimate biological fate of a system, the measurements are more complex and require specialized approaches for their analyses. Unfortunately, a majority of existing methods for RNA-seq data analysis treat gene expressions at sequential time points as repeated measurements and test DGE between conditions disregarding the ordinal aspect. They thus fail to highlight how genes exhibit a cascade of expression profiles over time.

### Modelling time course measurements

We demonstrate the utility of EMOGEA by applying it to a well-designed zebrafish embryogenesis data set (see Methods) previously generated and described by White and collegues^24^. Here the authors monitored mRNA expression of the developing zebrafish at 18 time points, with 5 replicate measurements at each time, covering 8 developmental stages. We analyzed the data by specifying 3, 4, 5 and 6 components (waves) which we modelled, extracted temporal profiles and evaluated them for consistency by determining (visually) if they differed sufficiently between each other. This type of sequential analysis is necessary because it is not possible to know, *a priori*, how many underlying expression profiles (waves) would be present in a data set, but alternative approaches exist for determining the number of components^25^. In principle, one can add as many components as there are samples, but degenerate solutions begin to emerge when the true number of unique profiles has been reached and, if an absurd number of profiles is specified, the resulting ill-conditioned matrix will be accompanied by non-convergence of the algorithm to which the user will be alerted.

The 6-component model of the zebrafish data and the extracted time profiles (normalized to unit length) are shown in Fig. 1b. These profiles are clearly very distinct from each other and represent groups of genes whose expression levels peak at different times during development. As an illustration, profiles I and II correspond to groups of genes at the very opposite ends of the gene expression spectra where one group peaks in expression towards the end of embryogenesis - in the larval stages (Profile II) - or is turned off soon after the embryos start development - at the cleavage stage (Profile I). In between these waves of gene expression, there are groups of genes whose expression levels peak at different developmental stages as shown in Profiles II to VI. Specifically, Profile III consists of genes peaking in expression during the early to mid-blastula stages, Profile IV and V consist of genes peaking at the late blastula and gastrula and, segmentation respectively. Finally, Profile VI consists of genes whose expression increases steadily with developmental time to pharyngula/hatching stage and then begins to slightly decrease in the final larval stage. We also show the empirical similarity between profiles modelled via EMOGEA and the expression levels of these groups of genes from the original data for genes correlated to Profiles I, V and VI in Fig. 1c.

**Figure 1:**
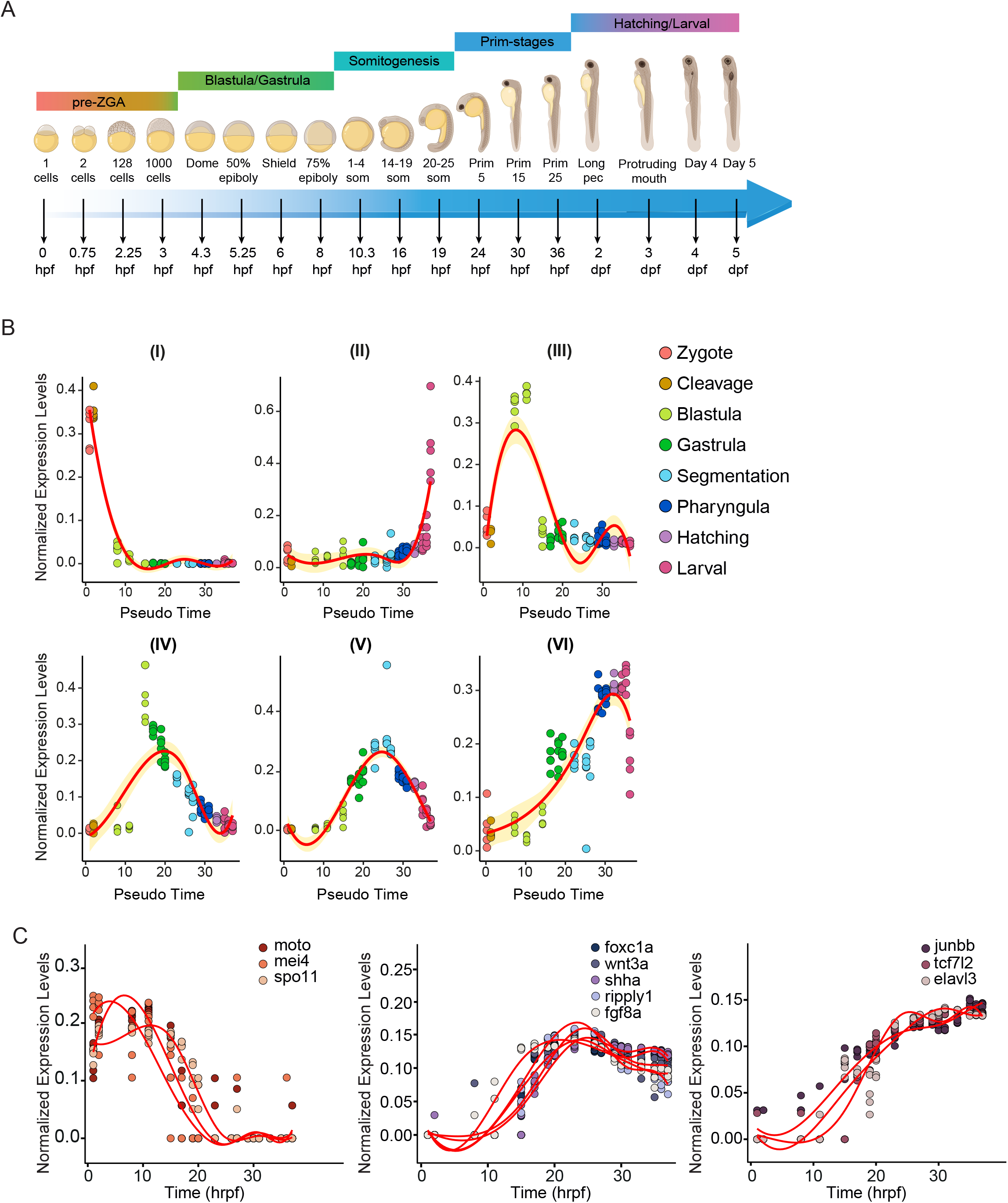
Results of the zebrafish embryogenesis RNA-seq data showing: (**A**) the developmental stages at which RNA-seq data were acquired spanning 0 hours post fertilization (hpf) to 5 days post-fertilization (dpf), and the embryonic stages represented by each time point; (**B**) the 6 profiles extracted by MCR via EMOGEA showing a cascade of temporal transcriptional events that peak at different critical developmental stages; (**C**) normalized expression levels of the genes whose temporal expression pattern best matches select profile I,V and VI is shown.

In order to more closely examine the groups of genes exhibiting specific characteristic waves of expression depicted in each of these six profiles, we calculated the cosine similarity between the expression of each gene, and each considered profile. We then ranked them in order of their strength of similarity (from 1 to 0) and took the top 200 genes whose temporal expression was most similar to each of the profiles. These genes are shown in Supplementary Table 1 for each profile. We further performed functional analyses in order to relate the gene expression levels to biological activity using over-representation analysis (ORA)^26^(Supplementary Fig. 1). This enabled us to objectively highlight characteristic functional pathways of specific stages of zebrafish embryogenesis.

Briefly, this analysis shows for instance, that Profile I comprises genes – such as *mei4*^27^, *moto*^28^ and *spo11*^29^ - involved in reproduction, gamete generation and germ cells development, consistent with their expression profile peaking in the zebrafish egg and zygote. Conversely, genes in Profile II are mainly *cyp* genes and *sult* superfamily members known to be highly expressed in the developed intestine and involved in the response to xenobiotics which is unsurprising since, at this stage, the zebrafish are free swimming and are introduced to environmental pollutants that require biotransformation to compounds that can be excreted more readily. Profile III chronologically follows Profile I and is characterized by intense cell replication (GO terms: chromosome segregation and reorganization), while Profile IV models the expression pattern of genes involved in endoderm development and embryo regionalization. This continues in Profile V with genes initiating segmentation and hatching. In the final developmental stages of the zebrafish (Profile VI) the peaking genes are those involved in extracellular structure organization (*i*.*e*. muscle development, cardiomyocyte differentiation).

The results show that EMOGEA yields gene expression profiles that represent groups of genes with similar modulations in their expression during embryogenesis. The relevance of the group of genes comprised in these profiles is supported by the biological functional analysis via ORA. This is an important and unique result that, unfortunately, cannot be obtained by methods that treat TC data as repeat measurements and analyze them for DGE. EMOGEA profiles highlight with clarity how the expression of different genes is modulated over time with a tractable biological interpretation. Although other methods such as DESeq2 have been extended to analyze time series data, they are still fundamentally based on pairwise comparisons, choosing for example a time point against which all other measurements are made.

### Modelling scRNA-seq data

Whereas, in principle, scRNA-seq data follow the design paradigm of case-control studies, they exhibit an inherent ordinal structure because (unless under exceptional circumstances) cells are not typically arrested at any stage of development, cell cycle or other transitions prior to sampling. Gene expression subsequently reflects this ordinal, cellular characteristic and there are several methods for ordering cells in a “pseudo time” to reflect this inherent structure. We used EMOGEA to model these data to reveal transcriptional profiles similar to those of bulk RNA-seq. To illustrate this utility, we use scRNA-seq data from Deng *et al*.,^30^ (see Methods), which were acquired to investigate allele-specific gene expression from cells dissociated from *in vivo* embryos of mouse pre-implantation covering 10 developmental stages from oocyte to blastocyst.

To employ our approach, we start by ordering the cells in pseudo time which we achieve by first performing principal component analysis (PCA) and plotting the data projections in first two principal components (PCs). We then determine the principal curve through the data cloud and obtain the orthogonal projections of the data onto this principal curve. We subsequently order the cells according to the principal curve projections as shown in Supplementary Fig. 2. Following this pseudo-temporal ordering, we analyzed the data by applying EMOGEA following a similar protocol to that used in bulk TC RNA-seq and show the 4 most distinctive expression profiles in Fig. 2b.

**Figure 2:**
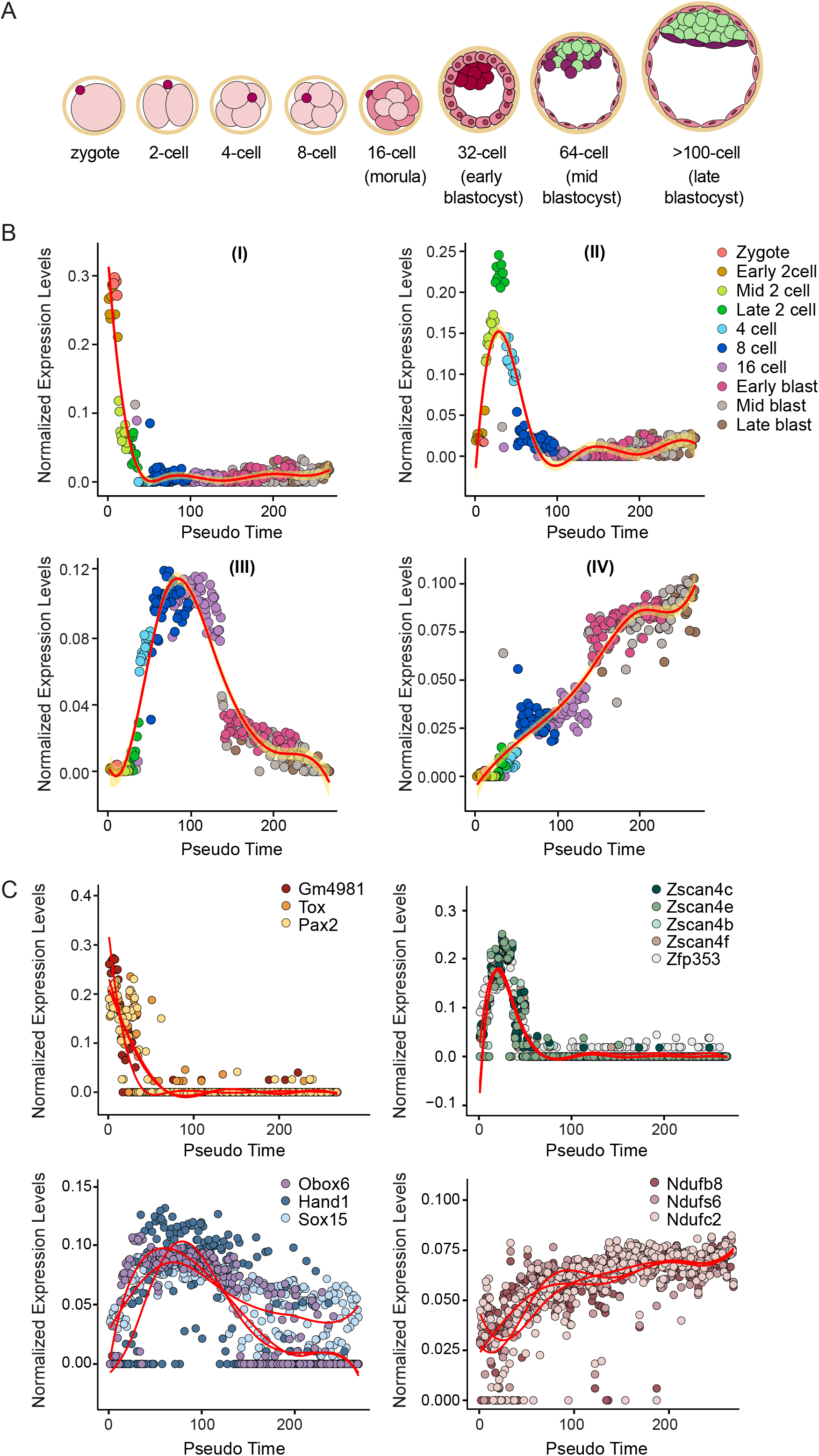
Results of scRNA-seq data analysis showing: (**A**) the developmental stages at which scRNA-seq data were acquired from zygotic embryo to blastocyst stages; (**B**) the 4 profiles extracted by MCR via EMOGEA showing a cascade of gene expression waves whose peaks at different developmental stages. The profile in panel (I) represents genes that have high expression in the zygotic stage but decrease in expression to baseline starting at the 4-cell stage. Panel (IV) shows genes with the opposite profile where expression starts out at baseline but increases to a maximum in the late blastocyst stage; (**C**), normalized expression levels of the genes whose temporal expression pattern best matches select profiles is shown in panels (I) to (IV) and whose function in described in the text.

The profiles reveal genes that are well organized in their expression levels to coincide with the emergence of new developmental stages. Particularly striking is the expression of genes shown in Profile I, which are turned off when the mouse embryos reach the 4-cell stage and are essentially not expressed throughout its development. Profile II, on the other hand, distinguishes genes that are expressed only at mid 2-cell stage, peak at the late 2-cell stage and begin to decrease their levels of expression in the late 4-cell and are back down to baseline expression at the 8-cell stage and onwards. What is interesting is that these genes are also not expressed at the zygote and early 2-cell stages. Profile III comprises genes that are expressed between the 4-cell and 16-cell stages while Profile IV consists of the genes whose expression rises steadily as the embryos develop.

To put the gene expression profiles into a context of mouse developmental biology, we selected some of the top genes in each profile (Supplementary Table 2), plotted their expression individually (Fig. 3c, and Supplementary Fig. 2b-c), and performed a cursory literature survey. The top gene in the list corresponding to Profile I, *Gm4981*, is reported^31^ to be part of the *Dux* locus which is structured as multiple repeats on the chromosome 10. This locus generates the earliest transcripts in the fertilized oocyte that subsequently initiate the transcription of the parental genomes, an event referred to as the embryonic genome activation (EGA) program^31,32^. EGA is a significant, temporally sensitive phase in normal mouse embryonic development and transcriptional events are well orchestrated. This is demonstrated by the expression of *Gm4981* which peaks immediately after fertilization of the oocyte (Profile I) to initiate the EGA program where target genes such as members of the *Zscan4* family, *Gm* family and *Tdpoz* family are highly transcribed (Profile II). Consistently, defects in the modulation of this critical phase of development have been correlated to phenotype defects of the morula and blastocyst stage. In subsequent profiles, we see that the temporal expression levels of *Obox6, Sox15* and *Hand1* show similarity to the trajectory of Profile III as they peak after the 4-cell state. First, Obox6 is a member of the Obox family of transcription factors and has recently been reported^33,34^ to be preferentially expressed in the oocyte, zygote, early embryos, and embryonic stem cells, where it regulates pluripotent stem cell reprogramming. Second, Sox15 is a member of the SRY-related HMG-box (Sox) family of transcription factors which is highly expressed in mouse undifferentiated embryonic stem cells and is progressively repressed upon cell differentiation. Interestingly, Sox15 can interact with different protein partners, such as Pou5F1 (known as Oct3/4) and Fhl3 (four and a half LIM domains 3) to regulate the Foxk1 gene (forkhead box protein K1), which is essential for the cell cycle progression of myogenic progenitor cells^35,36^. Finally, Sox15 is also involved in the differentiation of the trophoblast giant cell by enhancing the transcriptional activity of Hand1^37^, which is one of the top genes of Profile III. Profile IV mainly includes genes involved in metabolic processes, which become predominant in the final stages of organism development and in the adult animal.

**Figure 3:**
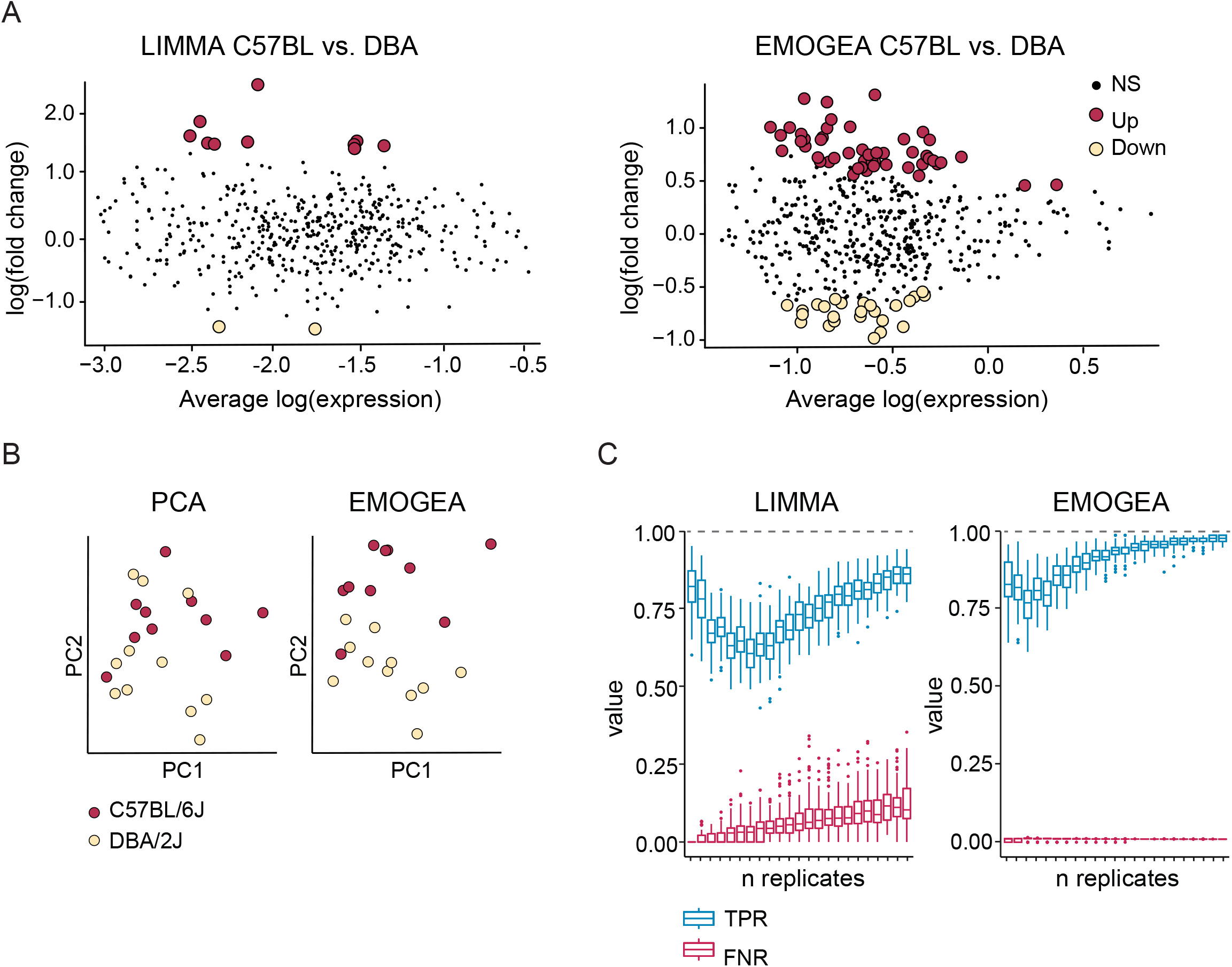
Analysis of case control studies showing: (**A**) differential expression analysis of least variable genes from *Bottomly* data set using limma before and after error weighting via EMOGEA, with similar adjusted \textit{p-value} cutoffs for differential expression; (**B**) exploratory analysis via PCA and EMOGEA of the least variable genes indicating that, without error weighting, it is not possible to distinguish the two mouse strains from transcripts with low expression levels and; (**C**) bootstrap estimates of the true positive rate (TPR) and true negative rate (TNR) for DGEA of the *Gerlinski* data using limma with and without error weighting via EMOGEA.

Examples include members of the NADH dehydrogenase [ubiquinone] family (*Nduf*) which form the NADH dehydrogenase complex I of the mitochondrial electron transport chain, glyceraldehyde-3-phosphate dehydrogenase (*Gapdh*), and *Hexa*, the alpha subunit of the lysosomal enzyme beta-hexosaminidase. These findings are consistent from the point of view of the animal’s increased energy demands as it develops, which subsequently lead to up-regulated expression of genes involved in energy metabolism.

### Modelling ‘case-control’ measurements

Case-control studies are by far the most common study designs in omics measurements. To demonstrate the importance of EMOGEA for analyzing data from such studies we employ a common data set first published by Bottomly and collegues^38^ in which expression levels of over 36,000 genes from the striatum of two mouse strains were measured. After error weighting via EMOGEA, we show results from three types of analyses in Fig. 3.

The first analysis represents differential expression analysis that focuses on low count signals. We selected 500 least variable genes (corresponding to low count signals) and performed DGEA on their expression levels using limma’s empirical Bayes moderated t-statistic for the two contrasts under the null hypothesis, before and after error weighting through EMOGEA. The mean-difference plot is shown in Fig 3a, and evidently illustrates that at the lower levels of gene count, the number of genes determined to be differentially expressed is significantly smaller without error weighting. The genes called differentially expressed by both approaches are shown in Supplementary Table 3 and highlight an important finding. Whereas analysis of the original data identified 1718 differentially expressed genes, our analysis uncovered an additional 54 differentially expressed genes that had been disregarded due to their low count signals. This finding is particularly significant because in many biological applications, low count signals associated with signaling genes and non-coding RNAs can have important biological implications, but are often not detected using available methods for DGEA.

The second analysis represents a popular approach to evaluating the results of a Case-Control experiment through exploratory analysis via methods such as principal component analysis (PCA), hierarchical clustering and multi-dimensional scaling (MDS) among others. Here we show, graphically, the separation between the two mice strains based on the expression levels of the least variable genes using PCA. In Fig. 3b, whereas the separation between the mouse strains is clear along PC2 from the EMOGEA analysis, this picture is less clear when measurement errors are not incorporated into the analysis. In many analyses PCA is performed either on the entire data set or on a pre-selected set of highly variable genes, an approach that implicitly focuses on the most dominant signals at the expense of the low intensity ones.

Finally, the third analysis focuses on perhaps one of the most important but often overlooked aspect of DGEA which is the rate of true positive and false negative discoveries. These two parameters are highly influenced by the level of uncertainty in the measurements as has been shown in Schurch *et al*.,^21^ and Gierlinski *et al*.,^39^. We sought to show the impact of measurement error weighting on these parameters using yeast RNA-seq data derived from two conditions with up to 48 replicates per experiment. We refer to this data set as the *Gerlinski* data which has been described extensively in the literature^21,39^. The data are particularly useful for demonstrating the usefulness of EMOGEA given its extensive unprecedented level of replication. Once again, we focus on low count signals to show the effectiveness of determining DGE from such measurements by generating from the original data, a pseudo-set of genes that are artificially “differentially expressed”. We achieved this by selecting from the expression measurements of wildtype samples, a set of 1000 genes with the lowest count levels. We subsequently selected form these, another 10% of the genes and inflated their counts to levels that exceeded the 95% limit of the data spread. This allowed us to obtain a dummy set of data that exhibited identical distributional characteristics to the original except for a known number of genes that were differentially expressed. We then analyzed these data using limma’s empirical Bayes moderated t-statistic, before and after processing with EMOGEA, with the objective of recovering the artificially differentially expressed genes. To avoid bias, we generated 1000 dummy sets of these “differentially expressed genes” via bootstrapping low count genes from the original data and calculated the true positive rate (TPR) and false negative rate (FNR) following the approach proposed by Schurch and collegues^21^. TPR corresponds to the ratio of genes determined to be differentially expressed to the total number of differentially expressed genes while FNR is the ratio of genes determined not to be differentially expressed to the total number of differentially ex-pressed genes.

## DISCUSSION

These results demonstrate several factors. First, analysis of bulk time course RNA-seq data of zebrafish embryogenesis using the EMOGEA framework display clear well-orchestrated waves of gene modulation that recapitulate the biological cascades expected during development. We show via functional analysis of the genes that correspond to each wave, the biological relevance of each profile and select a few genes that correlate with each profile to demonstrate their transcriptional trajectories.

Second, for scRNA-seq data we show again that this framework places the development of mouse embryos in the context of peaking gene expression at specific time points. With supporting biological inference from the literature, we show different sets of genes peaking in expression with a high degree of resolution to distinguish the zygotic, early, mid and late two-cell stages from each other. We use the method of principal curves to obtain a pseudo-temporal order and place the cells into their respective developmental continuum prior to applying multivariate curve resolution. Other methods for pseudo-time estimation are possible and we anticipate that this approach will be used to infer other non-branching cellular transitions exhibited by scRNA-seq experiments such as cell cycle, stem-cell trans-differentiation during organogenesis, T-cell activation among others.

Finally in relation to case-control studies we examine the question of highly variable (and therefore high count) genes and their influence on the results of DGE. Using a common data set, we show that the EMOGEA framework allows low count genes to model the biological question while approaches that ignore measurement error information are unable to differential expression from these low count signals. This has important consequences for studies involving non-coding RNAs and signaling genes that are generally expressed in low levels but have serious biological ramifications if they are differentially expressed.

Moreover, we employ the concept of true positive and negative rates to illustrate the effect or error weighting on these parameters. Schurch *et al*.,^21^ have shown that TPR should increase with the increasing number of replicates while the reverse is the case for the FNR. In. Fig 3c we show that the TPR after EMOGEA error weighting reaches a mean TPR of 90% after 8 replicates and stabilizes thereafter, while DGEA obtained by limma alone reaches a mean TPR of 63% after 8 replicates and does not stabilize until after 12 replicates and, at a mean value of only 72%. Even more striking is the FNR which stabilizes to 1% for EMOGEA analysis after 3 replicates. Surprisingly, the FNR for limma analysis alone starts out at 2% but rises as the number of replicates increases up to 4%. Over all the bootstrap experiments, EMOGEA results appear to be much more consistent looking at the spread of the bootstrap TPR and FNR values. These result are reassuring because they are consistent with the observations we made when analyzing the *Bottomly* data set^38^ where EMOGEA identifies more differentially expressed genes in the low gene-count range.

## METHODS

### Practical Implementation and Data Sets

#### Analysis of Zebrafish time course (TC) data

Zebrafish data from White *et. al*.,^24^ were used to demonstrate the utility of EMOGEA for time course data analysis. The data consist of poly(A) RNA expression profiles of baseline zebrafish embryogenesis capturing 18 time points covering 5 developmental processes. These were divided as: pre- and zygotic onset (covered by 4 time points); blastula/gastrula (covered by 4 time points); somitogenesis and prim (each covered by 3 time points) and; hatching/larval stages (covered by 4 time points). At each time point, 5 biological replicates of cDNA libraries were prepared from a pool of 12 embryos and sequenced on Illumina HiSeq 2500. The resulting sequences were aligned to the GRCz10 reference genome using TopHat2^40^ while gene counts were produced using HTSeq^41^.

We utilized data available as the Supplementary files in White *et al*.,^24^, which comprised of pre-processed and quality checked data consisting of 32,110 gene transcripts for each of the 90 cDNA libraries sequenced. Error information was obtained from the replicates associated with each time point and applied EMOGEA-MCR with weighting, specifying an incremental number of profiles to be modelled. We determined that 6 profiles sufficiently modelled the data in a way that we could see significant difference in the modulation of gene expression in line with developmental time-points and stages.

In order to determine which genes best matched the profiles, we computed a cosine similarity metric to assign a score between 0 and 1 (where a score of 1 is absolute similarity and 0 absolute dis-similarity) to identify which genes had expression profiles that were modulated in a similar temporal cascade as the EMOGEA profiles. This score was calculated as: 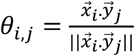, where 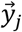 is the vector of expression profiles for gene *j* while 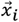 is profile *i*. It corresponds to the cosine of the angle between the two vectors and indicates the similarity of their approximate directionality. We then sorted the genes according to their (dis)similarities to the EMOGEA profiles and performed functional analyses on the topmost similar genes for each profile.

#### Analysis of single cell RNA-seq data

Single cell (sc)RNA-seq data utilized in this manuscript were first described in Deng *et al*.,^30^ from a study in which the authors sought to understand the genome wide allelic expression patterns in single cells obtained from two strains of mouse pre-implantation embryos. These data are particularly interesting in demonstrating the utility of EMOGEA, not only because of their temporal design, but also because scRNA-seq data display inherent ordinal characteristics. F1 embryos derived from 4 to 8 week-old CAST/EiJ mice mated with C57BL/6J mice were obtained from which 268 individual cells were dissociated at pre-implantation developmental stages ranging from oocyte to blastocyst. The single cells were then transferred into hypotonic Smart-seq lysis buffer and subjected to the Smart-seq2 protocol for the generation of RNA-seq libraries. Single-end sequencing at 46 to 59 bp of these libraries was then carried out on an Illumina HiSeq 2000. These data are available at the NCBI Gene Expression Omnibus (GEO, GSE45719). We accessed these data from GEO and mapped them to mm10 genome using STAR^42^ to yield a count matrix of dimension 22,431 transcripts x 268 cells. We subsequently estimated the pseudo-temporal order of the cells using principal curve and re-ordered each cell based on this order as shown in Supplementary Fig.1a. EMOGEA-MCR was used to obtain temporal profiles without weighting given that there were no replicate single cell data for this study. We similarly specified the number of profiles to be extracted incrementally and determined, for illustration purposes, that 4 profiles best described the developmental stages. Functional analysis was performed for genes whose expression profiles matched EMOGEA profiles as described for the bulk TC data.

### Analysis of Case-Control (CC) RNA-seq data

We utilized two data sets to demonstrate EMOGEA applications to CC studies. The first data set is the *Bottomly* data available at the NCBI Gene Expression Omnibus (GEO, GSE26024) consisting of RNA-seq measurements acquired to detect DGE between the striata of C56BL/6 and DBA/2J inbred mouse strains. The study included 21 samples (10 from C56BL/6 strain and 11 from DBA/2J strain) from which cDNA libraries were prepared and sequenced on an Illumina GAIIx generating an average of 22 million short sequencing reads. We aligned the reads to the mouse mm10 reference genome, generating a count matrix (using HTSeq^41^) of dimension 36,536 gene transcripts x 21 striata for both strains. In principle this corresponds to 10 biological replicates for the C56BL/6 strain and 11 biological replicates for the DBA/2J strain from which measurement errors could be calculated. Given the popularity of this data set, we used it to demonstrate the effects of EMOGEA analysis of the low intensity signals which were selected by choosing 1000 least variable genes.

The second data set is the *Gerlinski* data comprising of 48 replicate RNA-seq measurements of wild type (WT) *Saccharomyces cerevisiae* and corresponding snf2 (Δsnf2) knock-out mutant cell line. Poly(A) RNAs were extracted and converted to cDNA libraries that were sequenced (50bp single-end in hepta replicate), on an Illumina HiSeq 2000. We aligned the data obtained from the European Nucleotide Archive repository (ENA, PRJEB5348) to the Ensembl v64 release of the *S. cerevisiae* genome annotation with STAR^42^ and generated the count matrix using HTSeq^41^ to give a count matrix consisting of 7,076 gene features x 672 libraries sequenced (48 × 7 per strain).

### Theoretical Considerations (Online Methods)

Methods for the analysis of RNA-seq data operate on a “count” matrix ***X*** of dimension *n* x *m* where ***X***_*i j*_ is the number of reads assigned to transcript *i* in sequencing experiment *j*. Such matrices are produced by sequence alignment tools such as HTSeq^41^, featureCounts^43^, among others^13,44,45^. DGEA consists of: normalization of counts to remove systematic biases, estimation of parameters that describe the statistical model and, testing for differential expression or exploratory analysis using such methods as Principal Component Analysis (PCA) and Multi-Dimensional Scaling (MDS). Methods for gene expression analysis fall into categories distinguished by the statistic employed for testing differential expression. They include parametric t-test based methods *e*.*g*. Cuffdiff and Cuffdiff2^45,46^; and generalized least squares methods assuming Normal (after transformation), Poisson or negative binomial distributions *e*.*g*. edgeR, DESeq, DESeq2, baySeq, EBSeq, limma-voom^11,13,17,47–49^.

In this study we consider DGEA methods that utilize generalized linear models for parameter estimation to describe the relationships between gene expression and experimental conditions. We do not consider the importance of calculating gene counts but, rather focus on the importance of measurement errors in estimating these parameters - particularly for lowly expressed genes. Moreover, we do not engage in comparative analyses of these methods as there are numerous comprehensive reviews and extensive systematic comparisons between DGEA methods elsewhere with recommendations for best practices both for differential expression data pre-processing^13,39,50^.

Generalized linear models for DGEA are structured as follows. Considering a response *y*_*i*_ which might be an experimental condition such as disease, the expression of genes measured by RNA-seq for each patient can be represented as:

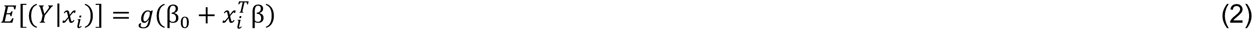

where E[Y] is the expected response, *g* the link-function, *β*_*0*_ the baseline, and *β* is a vector of parameters describing the relationship between the covariates, *x*_*i*_ (normalized gene counts) and the response. Typically, the variance of the expected values is represented as:

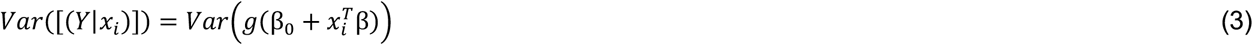

Numerous link functions can be used to describe the relationship between the response and predictors in *Eqn*. (2). Assuming identity link function for RNA-seq data, the maximum likelihood estimate of the parameters is obtained via ordinary least squares with an assumption that the variance, *Eqn*. (3), is identical, independent and normally distributed (*iid* normal) for all covariates. *Eqn*. (2) can subsequently be simplified as: *y* = *X*β, and the least squares estimate for the parameters given as: 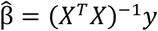. Zweiner *et al*.,^51^ have provided an expert evaluation of this, and other regression models, and shown that this solution is influenced by covariates that have high variance since they are not *iid* normal. In RNA-seq data, these covariates also correspond to gene transcripts with the high levels of expression. We have shown elsewhere^22^ using a simulated example that methods such as PCA also favor such covariates at the expense of low intensity ones. In that case we employed maximum likelihood PCA (MLPCA)^52^ to mitigate these effects by incorporating measurement errors during the course of estimating principal components with demonstrable success. We take the same approach here.

At its most basic form, MLPCA can be viewed as a superset of the classical PCA that is weighted by measurement errors that de-emphasize noisy measurements. It is however more sophisticated since it incorporates measurement errors of different structures ranging from the basic homoscedastic (*iid* normal) to more complex heteroscedastic noise with different correlation structures. Theoretical aspects of MLPCA are extensively covered in other references^52,53^ but we highlight two equations (*Eqns*. (5) and (6)) that show its fundamental differences and, hence, power. At the outset, it is important to declare that like PCA, MLPCA is a subspace estimation method that uses principles of maximum likelihood modelling to obtain a lower rank bilinear model for data in a high dimensional space. Subspace estimation methods reduce the dimensionality of data with a large number of features by transforming them to a new, a considerably smaller information-rich set that is devoid of noise. Using singular value decomposition (SVD) for example, a data matrix *X*_*m x n*_ can be represented as:

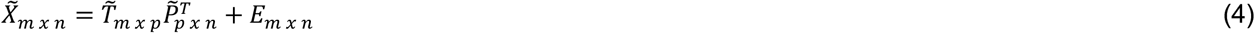

where 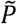 (loadings) describes the truncated set of new orthogonal axes and 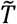, the sample coordinates in this new system. In conventional PCA, a new sample can be projected into the orthogonal subspace such that:

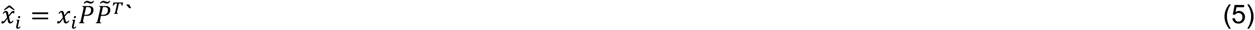

while the maximum likelihood estimate of *x*_*i*_ is given by a projection that is weighted by the errors in the measurements:

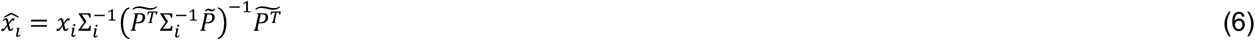

where Σ_*i*_ is the *n x n* error covariance matrix corresponding to sample *x*_*i*_. If the diagonal elements of Σ_*i*_ correspond to the variance of the features measured for sample *x*_*i*_ and the off-diagonals are all zeros, then *Eqn*. (6) is equivalent to *Eqn*. (5) and satisfies the assumptions of *iid* normal variance for the covariates. For many analytical measurements, especially RNA-seq, it is difficult to imagine a scenario where this would be true knowing that measurement errors are proportional to expression levels while the expression of genes is, in general, correlated.

#### Measurement Errors

Sources of variance in quantitative biology experiments in general can be represented in a compact way as:

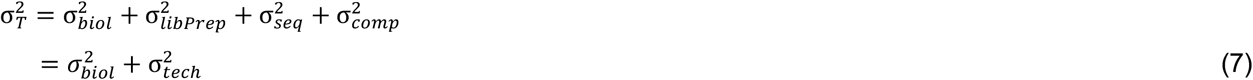

where 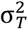 is the total variance, 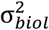 is the sample (biological) variance, 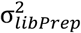 is the variance due to library preparation, 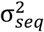 is the variance from sequencing and 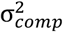 is the computational variance addressed by Pimentel *et al*.,^13^ and Robert *et al*.,^54^. The last three terms on the right-hand side of *Eqn*. (7) can be combined and referred to as the technical uncertainty and represented as 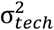. The magnitude of this term can be estimated through replication, and must be smaller than 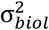 in order to test biological (rather than technical) hypotheses adequately as has been addressed in Conesa *et al*.,^55^. Using replicate measurements, 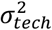 can be calculated for each sample as follows.

We assume that the mean of replicate measurements for each sample represents the true, *x*_*o*_, expression levels for each feature. Thus, the vector of measurement errors for each sample measurement, 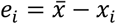, where 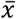 is the best estimate of *x*_*o*_ determined from replicate measurements. Of course, the more replicates one has the better the estimate. The error covariance matrix can then be used to characterize the statistical behavior of the vector of measurement errors and is defined as the outer product of the error vector such that:

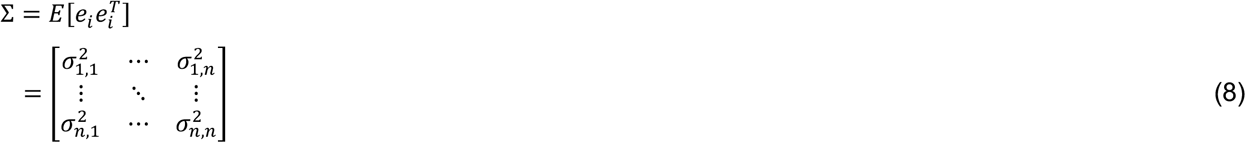

The diagonal elements of this matrix give the error variances associated with each feature and will therefore highlight any heteroscedasticity. For homoscedastic measurement errors, the off-diagonal elements, will be zeros (or approximately zero) while the values along the diagonals 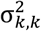 will be the same. Off-diagonal elements indicate the covariance of the measurement errors at features *j* and *k*.

#### Modelling Time Course Data with associated Measurement Errors

Given that many biological processes exist in a state of constant flux, time-ordered (ordinal) experiments provide key insights into the dynamics of cellular transitions as a result of: extraneous signals; developmental processes; and intrinsic, cyclic events such as cell cycle. Temporal RNA-seq data raise several experimental and computational challenges because the measurements exhibit complex properties that affect analysis and interpretations. Although many methods have been developed to analyze RNA-seq data from experiments designed under the ‘case-control’ setup, there have been relatively few computational developments for analyzing ordinal data. Most of the available methods perform a pairwise comparison of each time point to the first one, or to the same time point of a second time series/treatment, which ignores temporal dependencies and/biological insight that might propagate from one experimental time point to the next. There are several comparative studies^56^ for methods of analysis of time course data.

Our approach is conceptually simple and has been successfully used to model temporal DNA microarray data^57^ and metabolomics by both magnetic resonance and mass spectrometry^58^. We model RNA-seq data using the principles of bilinear modelling similar to *Eqn*. (4), with an approach that imposes alternative constraints to the solution of the first two lower rank matrices that comprise the right-hand side of the equation. This approach is unlike PCA which determines the solutions to *Eqn*. (4) by imposing the constraint that successive factors in the decomposition must (a) account for the largest amount of residual variance, and (b) be orthogonal to all of the factors determined to that point. Our approach, more generally referred to as multivariate curve resolution via alternating least squares (MCR-ALS), imposes a simple requirement of non-negativity in the elements of T and P. We and others^57–60^ have extensively covered the theoretical bases of MCR-ALS and it’s weighted alternative (MCR-wALS).

In brief, it is assumed that the expression matrix **X** can be decomposed into two linear matrices of lower rank, **T** and **P** similar to *Eqn*. (4). Without knowledge of either **T** or **P**, an initial guess of the expected number of components is made along with random positive numbers representing one of either 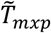, or 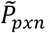.

Alternatively, a random set of vectors can be chosen from the count matrix, **X**, to represent 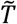 or 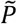. Once initial estimates are made, it is straightforward to determine the unknown via least squares, setting all negative values in the solution to zero. Suppose an initial estimate of 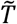 was made, the least squares estimate of 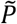 is simply 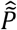, which is normalized to unit length and all its negative values set to zero. Subsequently using these values, 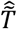 is estimated via least squares and its values constrained to be non-negative. This procedure is then repeated until some self-consistency criterion is met.

The weighted alternative to MCR is equally intuitive with the only addition being that measurement errors are incorporated in the estimation of matrices 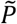 and 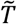. In cases where there was no measurement of gene expression (missing value) we assign an error of 9999, a disproportionately large error that downweighs the significance of the missing value. Considering the first half of the alternating LS procedure given **X**, which has an arbitrary error structure, and 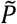, which is assumed to be known with certainty, we solve for 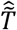. van Huffel *et al*.,^61^ show that the LS solution to 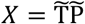 can be solved (conceptually) by first augmenting **X** with 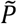 row-wise and finding the optimal *p*-dimensional subspace of the augmented matrix. In this case, it is clear that this subspace is defined by the *p* rows of 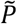, which are assumed to be known exactly. This problem can be simplified by determining the optimal representation of **X** in this subspace. Given measurement errors in **X** determined via replication, the estimate of **X** in the subspace of 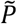 is then given by the maximum likelihood projection of **X** into the subspace of 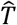 similar to *Eqn*. (6).

In the second half of the alternating LS procedure the estimate of **X** in the subspace of 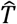 is determined via column wise maximum likelihood projection **X** into the space of 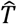 such that:

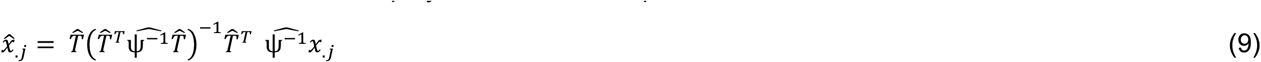

where 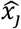 is the j^th^ column of **X** and *ψ* is the corresponding error covariance matrix, obtained as in *Eqn*. (7). The estimates for 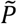 and 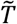 are determined, once again, via alternating LS using 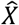 instead of **X**. As before, this process is repeated several times until a convergence criterion is minimized.

In this work, we set a maximum number of iterations to 200 while maximizing the self-consistency of \tilde{P} by minimizing the mean square error of estimation, that is, ((Σ(*P*^*new*^ *− P*^*old*^)^2^)/(*N* − 1)^1/2^ where P^old^ and P^new^ are the subsequent profile vectors, and *N* is the number of points that constitute 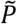. We then plot the vectors of **P** as a function of the ordinal variable to visualized how gene expression evolves over this variable.

### Graphical Output

Figure panels have been generated using Adobe Illustrator 22.1; scientific illustrations were created with the online web-based software BioRender (https://biorender.com/) and iStock (www.istockphoto.com/).

## Supplementary Figures

**Fig. S1**: Over Representation Analysis (ORA)^26^ was used to determine whether known biological processes were over-represented in the top 200 genes associated with each of the zebrafish embryogenesis profiles derived from EMOGEA. We show in bold-face representative biological processes for each profile.

**Fig. S2**: Panel (**A**) shows pseudo-temporal order of cells along the first principal curve with the position of the developmental trajectory occupied by each embryonic stage. Panel (**B**) shows the expression profile for a class of genes whose expression peaks between mid 2-cell and 4-cel stages similar to profile (II) in Figure 2. Panel (**C**) shows those genes whose expression continues to increase as the embryos develop from the zygotic to the late blastocyst stages similar to profile IV in Figure 2.

## Code And Data Availability

The source code and EMOGEA R package are available at: https://github.com/itikadi/EMOGEA. Specific code used to generate the results presented here and the processed data are available on Mendeley’s public data repository via this <u>link</u>.

## Author Contributions

T. K. K. Conceptualized the study, developed the models, analyzed all the data. T. K. K. and F.T. developed the R package and wrote the vignette. J.B. performed all the biological interpretation of the data, prepared and organized all Figures in the Manuscript. T. K. K. and J. B. wrote the manuscript. All authors discussed the results and the manuscript.

