## Supplementary figures and images for "ERROR MODELLED GENE EXPRESSION ANALYSIS (EMOGEA) PROVIDES A SUPERIOR OVERVIEW OF TIME COURSE RNA-SEQ MEASUREMENTS AND LOW COUNT GENE EXPRESSION"

### Supplemantary Figure 1

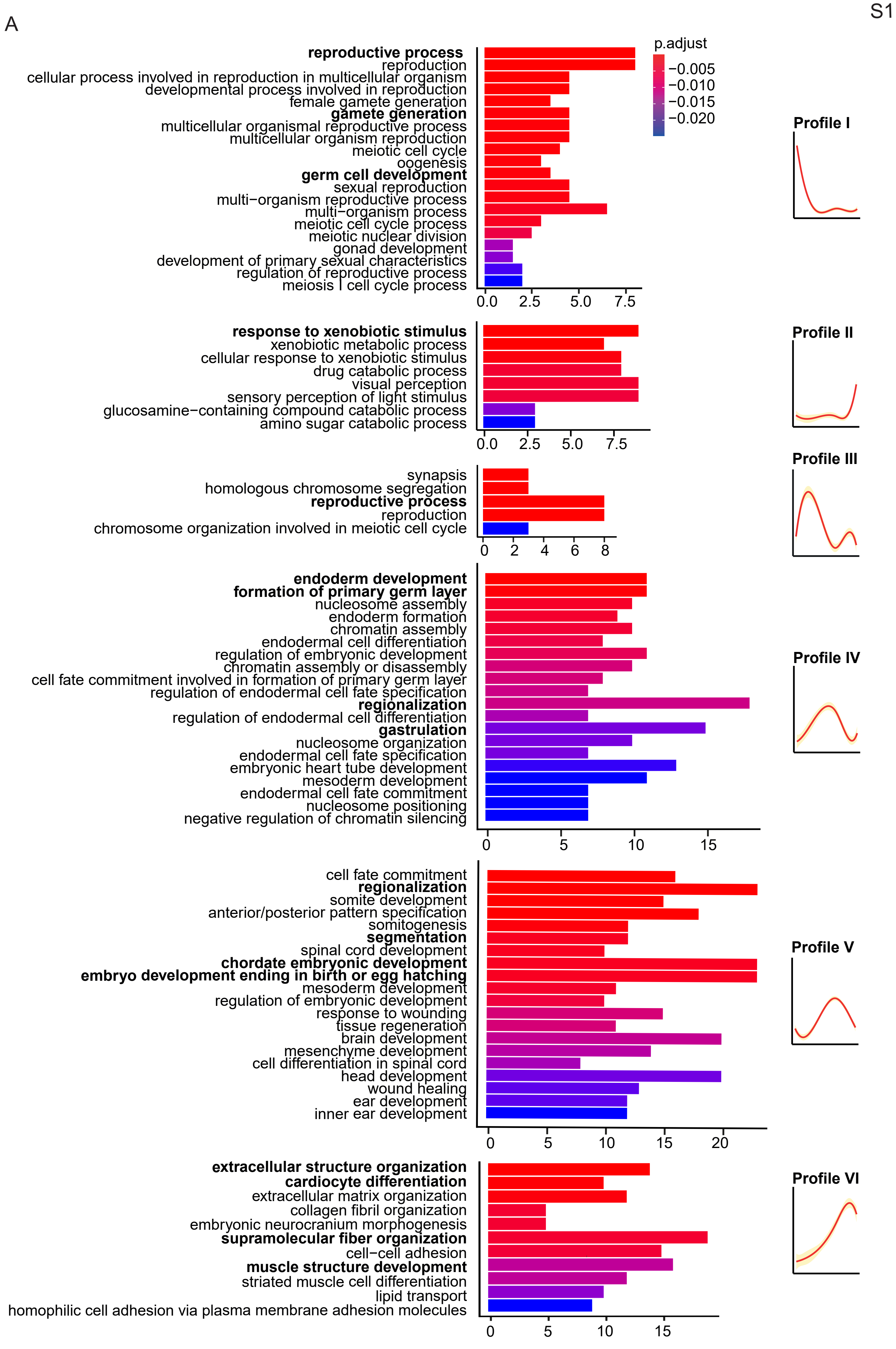

### Supplemental Figure 3

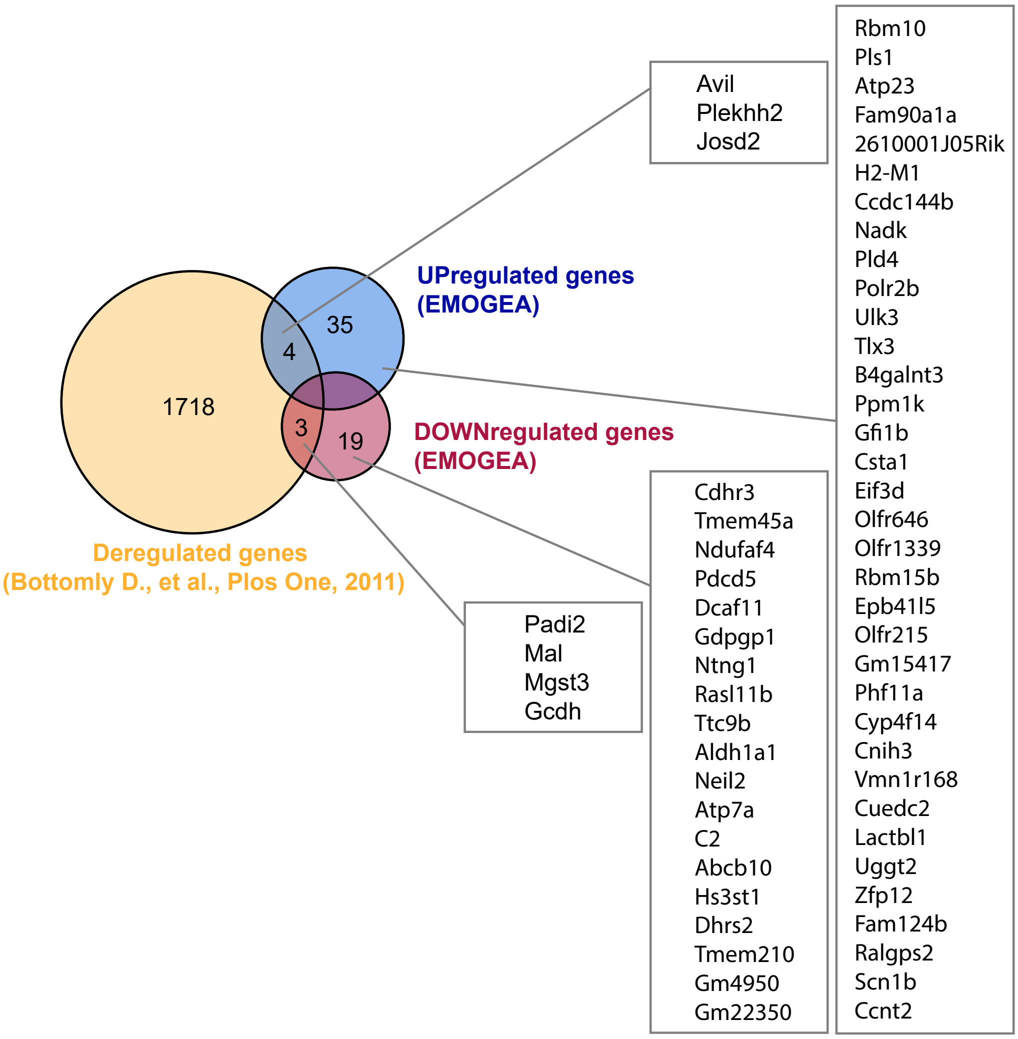

### Supplementary Figure 2

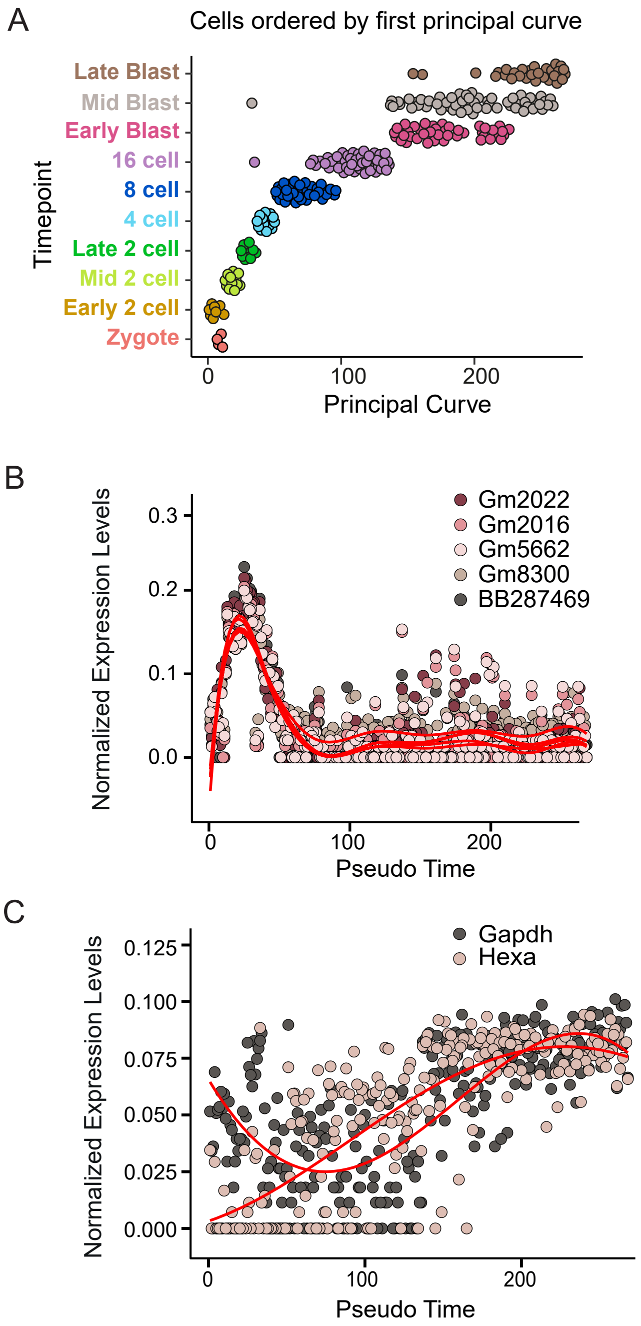
